# Proximity Biotin Labeling Reveals KSHV Interferon Regulatory Factor Networks

**DOI:** 10.1101/2020.10.16.343483

**Authors:** Ashish Kumar, Michelle Salemi, Resham Bhullar, Sara Guevara-Plunkett, Yuanzhi Lyu, Kang-Hsin Wang, Chie Izumiya, Mel Campbell, Ken-ichi Nakajima, Yoshihiro Izumiya

**Affiliations:** Department of Dermatology School of Medicine, University of California Davis (UC Davis), Sacramento, California USA; Genome Center, Proteomics Core, Genome and Biomedical Sciences Facility, UC Davis, Davis, California, USA; Department of Biochemistry and Molecular Medicine, School of Medicine, UC Davis, Sacramento, California USA; Viral Oncology and Pathogens-Associated Malignancies Initiative, UC Davis Comprehensive Cancer Center, Sacramento, California USA

**Keywords:** KSHV, TurboID, proteomics

## Abstract

Studies on “HIT&RUN” effects by viral protein are difficult when using traditional affinity precipitation-based techniques under dynamic conditions, because only proteins interacting at a specific instance in time can be precipitated by affinity purification. Recent advances in proximity labeling (PL) have enabled study of both static and dynamic protein-protein interactions. Here we applied PL method with recombinant Kaposi’s sarcoma-associated herpesvirus (KSHV). KSHV, a gamma-herpesvirus, uniquely encodes four interferon regulatory factors (IRFs 1-4) in the genome, and we identified KSHV vIRF-1 and vIRF-4 interacting proteins during reactivation. Fusion of mini-TurboID with vIRF-1 or vIRF-4 did not interfere with KSHV gene expression, DNA replication, or *de novo* infections. PL identified 213 and 70 proteins for vIRF-1 and vIRF-4 respectively, which possibly interact during KSHV reactivation, and 47 of those were shared between the two vIRFs; the list also includes three viral proteins, ORF17, thymidine kinase, and vIRF-4. Functional annotation of respective interacting proteins showed highly overlapping biological functions such as mRNA processing and transcriptional regulation by TP53. Involvement of commonly interacting 44 cellular proteins in innate immune regulation were examined by siRNAs, and we identified that splicing factor 3B (SF3B) family proteins were clearly involved in interferons transcription and suppressed KSHV reactivation. We propose that recombinant TurboID-KSHV is a powerful tool to probe key cellular proteins that play a role in KSHV replication, and selective splicing factors may have a function beyond connecting two exon sequences to regulate innate immune responses.

**Importance:** Viral protein interaction with a host protein shows at least two sides: (i) taking host protein functions for its own benefit and (ii) disruption of existing host protein complex formation to inhibit undesirable host responses. Due to use of affinity-precipitation approaches, the majority of our studies focused on how the virus takes advantage of the newly-formed protein interactions for its own replication. Proximity labeling (PL) however, can also highlight the transient and negative effects – those interactions which lead to dissociation from the existing protein complex. Here we highlight the power of PL in combination with recombinant KSHV to study viral host interactions.

## Introduction

Kaposi sarcoma herpesvirus (KSHV) is a pathogen associated with endothelial Kaposi’s sarcoma (KS) (1, 2), B-cell malignancies such as primary effusion lymphoma (PEL), and AIDS related multicentric Castleman’s disease (MCD) (3–6). In these cancer cells, KSHV mostly exhibits latent infection, where most of the viral genes are silenced to escape recognition by the host immune system. However, small population of infected cells undergo spontaneous reactivation, where all of the KSHV genes are expressed for production of progeny virions. Although lytic replication produces infectious virions and facilitates transmission of the virus to neighboring cells or host, it also increases the risk of the virus being caught by the host immune system (7). Host immune systems detect pathogens through binding of pathogen associated molecular pattern (PAMPs) to pattern recognition receptors (PRRs). Several PRRs such as IFI16 (8, 9), RIG-I (10–12), TLR9 (13), TLR3 (14), TLR4 (15), and NLRP1 (16) are known to detect KSHV associated PAMPs. The recognition of KSHV DNA by PRRs leads to phosphorylation, dimerization, and nuclear translocation of IRF3/IRF7. IRF3/IRF7 binds to DNA through its DNA binding domain (DBD), which results in secretion of cytokines and interferons (IFN). To counteract the host response, KSHV encodes several immunomodulatory proteins such as viral-interferon regulatory factors (vIRFs) that inhibit the antiviral response and aid viral replication (17, 18).

KSHV genome encodes four vIRFs, vIRF-1-4. The N-termini of vIRFs exhibit similarity to N-termini of cellular IRFs, however viral IRFs lack a key tryptophan residue, which is required for binding to DNA (19). vIRF-1, vIRF-2, and vIRF-4 are inducible lytic genes, although vIRF-1 can also be found in a small portion of latently infected cells. In contrast, vIRF-3 (also known as LANA2) was discovered as a latent protein and its expression remains unchanged during reactivation (20). Studies on the function of vIRFs found that vIRFs counteract the host IFN response by interacting with cellular proteins. vIRF-1 suppresses cellular IRF3-mediated transcription by binding to p300, thereby preventing p300/CBP-IRF3 complex formation (21, 22). vIRF-1 also promotes KSHV lytic replication by recruitment of USP7 (23). vIRF-2 was found to inhibit KSHV lytic gene expression by increasing the expression of cellular antiviral factors like Promyelocytic leukemia nuclear bodies (PML) (24). Similarly, vIRF-3 suppresses KSHV reactivation by interacting with USP7, and the interaction also supports PEL cell growth (23). Furthermore, vIRF-4 has been found to play a crucial role in triggering the KSHV latency-to-lytic switch through interfering with the BCL6-vIRF-4 axis (25). vIRF-4 also associates with IRF7, and inhibits IRF7 dimerization to suppress IFN production (26). These studies sometimes showed different results in different cell lines, suggesting the significance of implementing proteomic approaches that can reveal vIRFs interaction networks more comprehensively. A broader view of the vIRFs interactomes will certainly help to understand their diverse protein functions.

Dynamic and stable protein-protein interactions are key to cellular processes and biological pathways. Affinity purification coupled with mass spectrometry (AP-MS) has been an invaluable method used to identify protein-protein interactions. However, AP-MS often fails to identify weakly or transiently interacting proteins. To overcome this drawback, enzyme-based Proximity-based labeling (PL) approaches have been developed. The approach provides sensitivity and specificity required to study dynamic protein-protein interaction. BirA_R118G_ (BirID) was the first proximity-based labelling enzyme identified in *E.coli* which conjugates biotin to lysine residues of neighboring proteins (27). However, original BirID required the presence of biotin for several hours to be able to biotinylate a sufficient amount of proteins for analysis, thereby restricting its use for dynamic processes. Recently, two variants of BirID have been developed by directed evolution named as mini-TurboID (28 kD) and TurboID (35 kD), which allow proximity labeling in less than 10 min without significant toxicity (28). The TurboID based approach has already been successfully employed in a wide variety of species including mammalian cells (28–32), *Drosophila* (33), plants (34–37), yeast (38), flies and worms (28). In this study, we prepared recombinant 3xFlag-mini-TurboID-vIRF-1 and 3xFlag-mini-TurboID-vIRF-4 KSHV that employs mini-TurboID to biotinylate host and viral proteins in vicinity to these two viral proteins. The proximity-labeling approach combined with mass spectrometry identified both previously-identified cellular proteins, as well as new host proteins as their interacting partners. The siRNA screenings of these interacting proteins identified that selective splicing factors function to suppress KSHV reactivation and are associated with anti-viral responses.

## Materials and Methods

### Chemicals

Dulbecco’s modified minimal essential medium (DMEM), Fetal bovine serum (FBS), phosphate buffered saline (PBS), Trypsin-EDTA solution, 100x Penicillin-streptomycin-L-Glutamine solution and Strep-HRP conjugate were purchased from Thermo Fisher (Waltham, MA USA). Puromycin and G418 solution were obtained from InvivoGen (San Diego, CA, USA). Hygromycin B solution was purchased from Enzo Life Science (Farmingdale, NY, USA). Anti-ORF57, anti-K8, and anti-K8.1, antibodies were purchased from Santa Cruz Biotechnology Inc (Santa Cruz, CA, USA). Anti-K-Rta antibody was described previously (39). All other chemicals were purchased from Millipore-Sigma (St. Louis, MO, USA) unless otherwise stated.

### Cells, siRNA transfection and reagents

iSLK.219 cells were maintained in DMEM medium supplemented with 10% FBS, 10 μg/ml puromycin, 400 μg/ml hygromycin B, and 250 μg/ml G418. iSLK cells were maintained in DMEM medium supplemented with 10% FBS, 1% penicillin-streptomycin solution and 10 μg/ml puromycin. iSLK cells were obtained from Dr. Don Ganem (Novartis Institutes for Biomedical Research). A549 cells were obtained from Dr. Tsucano (University of California, Davis). A549 cells were grown in DMEM containing 10% FBS and 1% penicillin-streptomycin. Transfection of siRNA in iSLK.219 cells was performed with Lipofectamine RNAiMax reagent (Invitrogen) according to manufacturer’s protocol.

### Quantification of viral replication

siRNA targeting the cellular genes were transfected in iSLK.219 cells for 48h followed by KSHV reactivation by doxycycline (1 μg/ml). After 24h, the RFP fluorescence intensity was quantified using ImageJ software. The RFP signal intensity was normalized relative to non-targeting siRNA (NTC).

### Construction of vIRF-1 and vIRF-4 miniTurbo KSHV BAC16

Recombinant KSHV was prepared by following a protocol for En passant mutagenesis with a two-step markerless red recombination technique (40). Briefly, codon optimized mini-TurboID coding sequence (Table 1), which also encodes 3x Flag tag was first cloned into a pBS SK vector (Thermo Fisher, Waltham, MA USA). The pEPkan-S plasmid was used as a source of the kanamycin cassette, which includes the I-SecI restriction enzyme site at the 5’-end of kanamycin coding region (40). Kanamycin cassette was amplified with primer pairs listed in Table 1, and cloned into the mini-TurboID coding region at a unique restriction enzyme site. The resulting plasmid was used as a template for another round of PCR to prepare a transfer DNA fragment for markerless recombination with BAC16 (41). Recombinant BAC clones with insertion and also deletion of the kanamycin cassette in the BAC16 genome were confirmed by colony PCR with appropriate primer pairs. Recombination junctions and adjacent genomic regions were amplified by PCR and the resulting PCR products were directly sequenced with the same primers to confirm in-flame insertion of mini-TurboID cassette into the BAC DNA. The resulting recombinant BAC was confirmed by restriction enzyme digestions (*Hind*III and *Bgl*II), if there were any large DNA deletions. Two independent BAC clones were generated for each mini-TurboID tagged recombinant virus as biological replicates, and used one of the clone for protein ID. Entire BAC DNAs were subsequently sequenced.

**Table 1.**
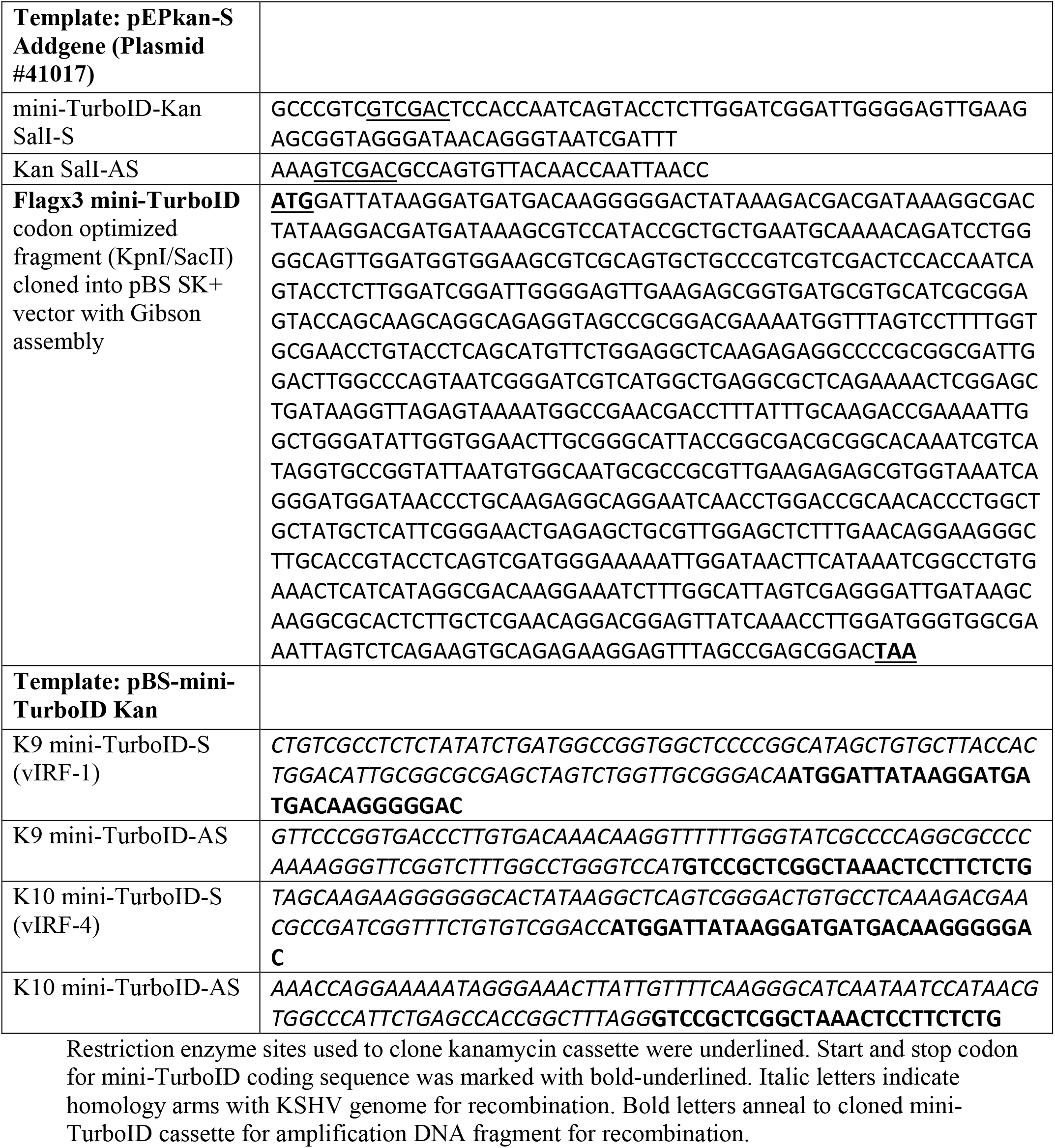
Primers, Plasmid, and Gene block DNA sequence used for BAC16 recombination (5’->3’)

### Western blotting

Cells were lysed in IP lysis buffer (25 mM Tris-HCl pH 7.4, 150 mM NaCl, 1% NP-40, 1 mM EDTA, 5% glycerol) containing protease inhibitors (Roche, Basel, Switzerland). Total cell lysates (25 μg) were boiled in SDS-PAGE loading buffer and subjected to SDS-PAGE and subsequently transferred to a polyvinylidene fluoride membrane (Millipore-Sigma, St. Louis, MO, USA) using a semidry transfer apparatus (Bio-Rad, Hercules, CA, USA). Streptavidin-HRP conjugate was used at 1:3000 dilution. Final dilution of the primary antibody was 1:5,000 for anti-K-Rta rabbit serum, 1 μg/mL of anti-K8α (Santa Cruz, Santa Cruz, CA, USA), 1 μg/mL of anti-ORF57 mouse monoclonal antibody (Santa Cruz, Santa Cruz, CA, USA), 1 μg/mL of anti-K8.1 mouse monoclonal (Santa Cruz, Santa Cruz, CA, USA), and 1:5,000 for anti-β-actin mouse monoclonal (Millipore-Sigma, St. Louis, MO, USA). Washing membranes and secondary antibody incubations were performed as described previously (42).

### Quantification of viral copy number

Two hundred microliter of cell culture supernatant was treated with 12 μg/ml of DNase I for 15 min at room temperature to degrade uncapsidated DNA. This reaction was stopped by the addition of EDTA to 5 mM followed by heating at 70°C for 15 min. Viral genomic DNA was purified using QIAamp DNA Mini Kit according to the manufacturer’s protocol, and eluted in 100 μl of buffer AE. Four microliters of eluate was used for real-time qPCR to determine viral copy number, as described previously (42).

### Preparation of purified KSHV

iSLK cells latently infected with mini-TurboID-KSHVs were seeded in eight to ten 15 cm dishes, and stimulated with 1 μg/mL of doxycycline and 3 mM sodium butyrate (NaB) for 24 h and further incubated with culture media without stimuli for 72 h. The culture supernatant was centrifuged using the Beckman SW28 rotor (25,000 rpm, for 2 h) with 25% sucrose cushion. Virus pellet was dissolved in DMEM and further purified by discontinuous sucrose gradient (25-60%) centrifugation using Beckman SW40Ti rotor (21,000 rpm, for 16 h). The virus pellet was dissolved in DMEM for infection.

### Real time RT-PCR

Total RNA was isolated using Quick-RNA miniprep kit (Zymo Research, Irvine, CA, USA). First strand cDNA was synthesized using High Capacity cDNA Reverse Transcription Kit (Thermo Fisher, Waltham, MA USA). Gene expression was analyzed by realtime qPCR using specific primers for KSHV ORFs designed by Fakhari and Dittmer (43). We used 18S ribosomal RNA as an internal standard to normalize viral gene expression.

### Affinity purification of biotinylated proteins

The affinity purification was done with streptavidin coated magnetic beads (Thermo-Fisher). Briefly, 150 μl magnetic beads/sample were pre-washed with RIPA lysis buffer (150 mM NaCl, 5 mM EDTA (pH 8), 50 mM Tris (pH 8), 1% NP-40, 0.5% sodium deoxycholate, 0.1% SDS) 3 times. Total 3 mg of whole cell lysate was incubated with pre-washed streptavidin beads at room temperature for 1h for rotation. The beads were collected using magnetic stand and washed three times with wash buffer according to manufacturer’s protocol. Finally, beads were resuspended in 200 μl of wash buffer and sent to UC Davis Proteomics core for on bead digestion and LC-MS/MS analysis.

### MS sample preparation

Protein samples on magnetic beads were washed four times with 200 μl of 50mM ammonium bicarbonate (AMBIC) with a twenty-minute shake time at 4°C in between each wash. Roughly 2.5 μg of trypsin was added to the bead and AMBIC and the samples were digested over night at 800 rpm shake speed. After overnight digestion, the supernatant was removed, and the beads were washed once with enough 50 mM ammonium bicarbonate to cover. After 20 minutes at a gentle shake the wash is removed and combined with the initial supernatant. The peptide extracts are reduced in volume by vacuum centrifugation and a small portion of the extract is used for fluorometric peptide quantification (Thermo scientific Pierce). One microgram of sample based on the fluorometric peptide assay was loaded for each LC-MS analysis.

Digested peptides were analyzed by LC-MS/MS on a Thermo Scientific Q Exactive Orbitrap Mass spectrometer in conjunction Proxeon Easy-nLC II HPLC (Thermo Scientific) and Proxeon nanospray source. The digested peptides were loaded a 100 micron × 25 mm Magic C18 100Å 5U reverse phase trap where they were desalted online before being separated using a 75 micron × 150 mm Magic C18 200Å 3U reverse phase column. Peptides were eluted using a 60-minute gradient with a flow rate of 300 nl/min. An MS survey scan was obtained for the m/z range 300-1600, MS/MS spectra were acquired using a top 15 method, where the top 15 ions in the MS spectra were subjected to HCD (High Energy Collisional Dissociation). An isolation mass window of 2.0 m/z was for the precursor ion selection, and normalized collision energy of 27% was used for fragmentation. A fifteen second duration was used for the dynamic exclusion.

### MS/MS analysis

Tandem mass spectra were extracted and charge state deconvoluted by Proteome Discoverer (Thermo Scientific) All MS/MS samples were analyzed using X! All MS/MS samples were analyzed using X! Tandem (The GPM, thegpm.org; version X! Tandem Alanine (2017.2.1.4)). X! Tandem was set up to search the Human and Kaposi Sarcoma Herpes virus database (149182 entries) assuming the digestion enzyme trypsin. X! Tandem was searched with a fragment ion mass tolerance of 20 PPM and a parent ion tolerance of 20 PPM. Carbamidomethyl of cysteine and selenocysteine was specified in X! Tandem as a fixed modification. Glu->pyro-Glu of the N-terminus, ammonia-loss of the N-terminus, gln->pyro-Glu of the N-terminus, deamidated of asparagine and glutamine, oxidation of methionine and tryptophan and dioxidation of methionine and tryptophan were specified in X! Tandem as variable modifications.

Scaffold (version Scaffold_4.8.4, Proteome Software Inc., Portland, OR) was used to validate MS/MS based peptide and protein identifications. Peptide identifications were accepted if they could be established at greater than 98.0% probability by the Scaffold Local FDR algorithm. Peptide identifications were also required to exceed specific database search engine thresholds. Protein identifications were accepted if they could be established at greater than 5.0% probability to achieve an FDR less than 5.0% and contained at least 2 identified peptides. Protein probabilities were assigned by the Protein Prophet algorithm (44). Proteins that contained similar peptides and could not be differentiated based on MS/MS analysis alone were grouped to satisfy the principles of parsimony. Proteins sharing significant peptide evidence were grouped into clusters.

### Pathway analysis

The proteins identified to be interacting with vIRF-1 and vIRF-4 were used for Gene ontology and network analysis. The top gene ontology processes were enriched by Metascape web-based platform, and the Metascape software was used for gene ontology and network analysis (45).

### Statistical analysis

Results are shown as mean ± SD from at least three independent experiments. Data was analyzed using unpaired Student’s t test, or ANOVA followed by Tukey’s HSD test. A value of p<0.05 was considered statistically significant.

## Results

### Construction of 3xFlag-mini-TurboID-K9 and 3xFlag-mini-TurboID-K10 KSHV BAC16

Biotin labelled proximity labeling (PL) has emerged as a powerful method for probing various target proteins in a wide variety of species including mammalian cells and unicellular organism (28, 32, 33, 35, 37, 38). We thought that applying the technique to virology would be particularly beneficial, because not only viral-host interactions are inherently dynamic but viruses are also completely dependent on host cell machinery for their replication. In fact, many key cellular proteins, such as p53, were identified from virology as viral protein interacting proteins. Our major goal is thus to report on the successful application and utility of PL in conjunction with recombinant KSHV BAC system.

To generate recombinant KSHV conveniently, we first prepared a template plasmid, which is used to create PCR fragments for recombination. The template encodes a 3xFlag tag at the N-terminus of mini-TurboID and kanamycin cassettes in mini-TurboID coding region as an excisable format with I-SecI induction. The 3xFlag-mini-TurboID-kana DNA fragment was amplified with primers with homology arms, and amplified fragments were then used for recombination by using a two-step recombination approach as previously described (40) (**Figure 1A**). The 3xFlag-mini-TurboID-K9 and 3xFlag-mini-TurboID-K10 BAC16 were directly transfected into iSLK cells and selected with hygromycin (1 mg/ml) to generate iSLK cells harboring latent 3xFlag-miniTurboID-K9 KSHV genome (named as vIRF-1 mini-TurboID cells) and 3xFlag-mini-TurboID-K10 KSHV (named as vIRF-4 mini-TurboID cells) (**Figure 1B**).

**Figure 1.**
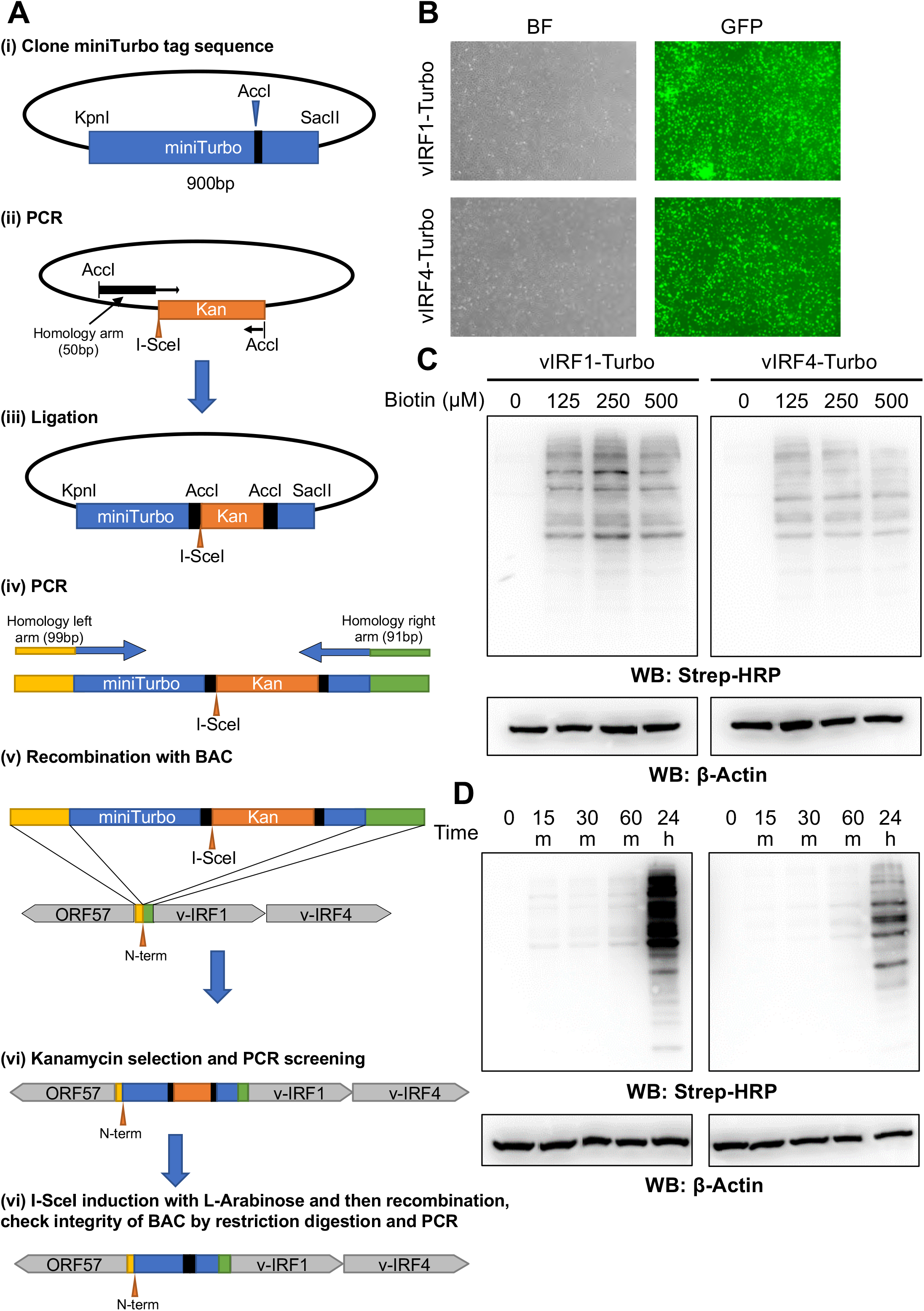
Engineering of mini-TurboID KSHVs. **(A) Schematic diagram for construction of 3xFlag-miniTurboID-K9 and 3xFlag-miniTurboID-K10 KSHV BAC16**. (i) The codon optimized cDNA fragment (900 bp) of mini-TurboID was synthesized and cloned into pBS vector between KpnI and SacII restriction enzyme sites. (ii) The kanamycin cassette with I-SceI recognition sequence along with 50 bp homologous sequence was generated by PCR with pEP-Kan plasmid as a template, and cloned into *Acc*I restriction enzyme site. (iii-v) The resulting plasmid was fully sequenced and used as a template to generate a DNA fragment for homologous recombination with BAC16 inside bacteria. (vi, vii) After confirmation of insertion at correct site by colony PCR screening, the kanamycin cassette was deleted by recombination with induction of I-SceI in bacteria by incubating with L-Arabinose. Correct insertion of the mini-TurboID and integrity of BAC DNA were confirmed by sequencing of PCR-amplified fragments and restriction digestions. Primers and DNA fragment used are listed in Table 1. **(B) Generation of vIRF-1 and vIRF-4 TurboID stable cells.** iSLK cells were transfected with 3xFlag-miniTurboID-K9 and 3xFlag-miniTurboID-K10 KSHV BAC16 and stably selected with hygromycin (1 mg/ml). GFP images show iSLK latently infected with 3xFlag-miniTurboID-K9 (upper panels) and 3xFlag-miniTurboID-K10 KSHV BAC16 (lower panels). BF: Bright Field, GFP: Green fluorescent protein. **(C) Biotin ligase activity of mini-TurboID tagged vIRF-1 and vIRF-4.** The vIRF-1 and vIRF-4 mini-TurboID cells were stimulated with Dox (1μg/ml) and NaB (3 mM) for 24h followed by incubation with indicated concentration of D-biotin for 1h. Activity of mini-TurboID was examined by immunoblot using Streptavidin HRP conjugate. WB: Western Blot. **(D) Dependency of mini-TurboID on labelling time.** vIRF-1-Turbo and vIRF-4 mini-TurboID cells were stimulated with Dox (1 μg/ml) and NaB (3 mM) for 24 h followed by incubation with D-biotin (500 μM) for indicated time-points. Activity of mini-TurboID was analyzed by immunoblot using Streptavidin HRP conjugate. WB: Western Blot, m: minutes, h: hours.

Next, optimal concentration of exogenous biotin and duration of incubation time for efficient labelling was determined. The vIRF-1 and vIRF-4 mini-TurboID cells were reactivated for 24 h, incubated with varying concentration of biotin (0, 125, 250 and 500 μM) for 1 h, and subsequently monitored for their biotinylation signal in whole cell lysates using streptavidin immunoblots. Untreated cells in absence of biotin were used as negative control. Immunoblot analysis using the HRP-conjugated streptavidin showed multiple biotinylated protein indicating successful labelling of proteins with vIRF-1 and vIRF-4 tagged mini-TurboID. Comparable levels of signal intensity were observed until 500 μM, suggesting that saturation of protein biotinylation occurs at 125 μM (**Figure 1C**). Similarly, vIRF-1 Turbo and vIRF-4 mini-TurboID cells were incubated with biotin for various time periods. Biotinylation signal was seen within 15 mins after addition of exogenous biotin and streptavidin signals gradually increased along with incubation time (**Figure 1D**). Considering only a small proportion of cells were reactivating in a dish, we concluded that there was a sufficient amount of biotinylation in the cells for protein identification. For following studies, we decided to use a saturating amount of biotin (500 μM) for 60 mins incubation.

### Gene expression in vIRF-1 and vIRF-4 mini-TurboID cells

We next verified the induction of viral genes to ensure that tagging K9 or K10 gene with 3xFlag-mini-TurboID has little effects on viral gene expression and replication. For this, we stimulated vIRF-1 and vIRF-4 mini-TurboID cells with Doxycycline (Dox) and performed qPCR for selected KSHV genes. We observed induction of KSHV lytic genes, PAN RNA, ORF6, vIRF-1 and vIRF-4 (**Figure 2A**). In addition, we verified induction of selected lytic KSHV proteins at 48h (**Figure 2B**), and virion production in culture supernatant at 96 h post-reactivation. Finally, culture supernatant was also used to infect A549 recipient cells to verify infectivity (**Figure 2C**). Altogether, these observations indicate that 3xFlag-mini-TurboID protein tag did not interfere with viral gene expression and that the recombinant KSHVs are replicating to produce infectious viral particles.

**Figure 2.**
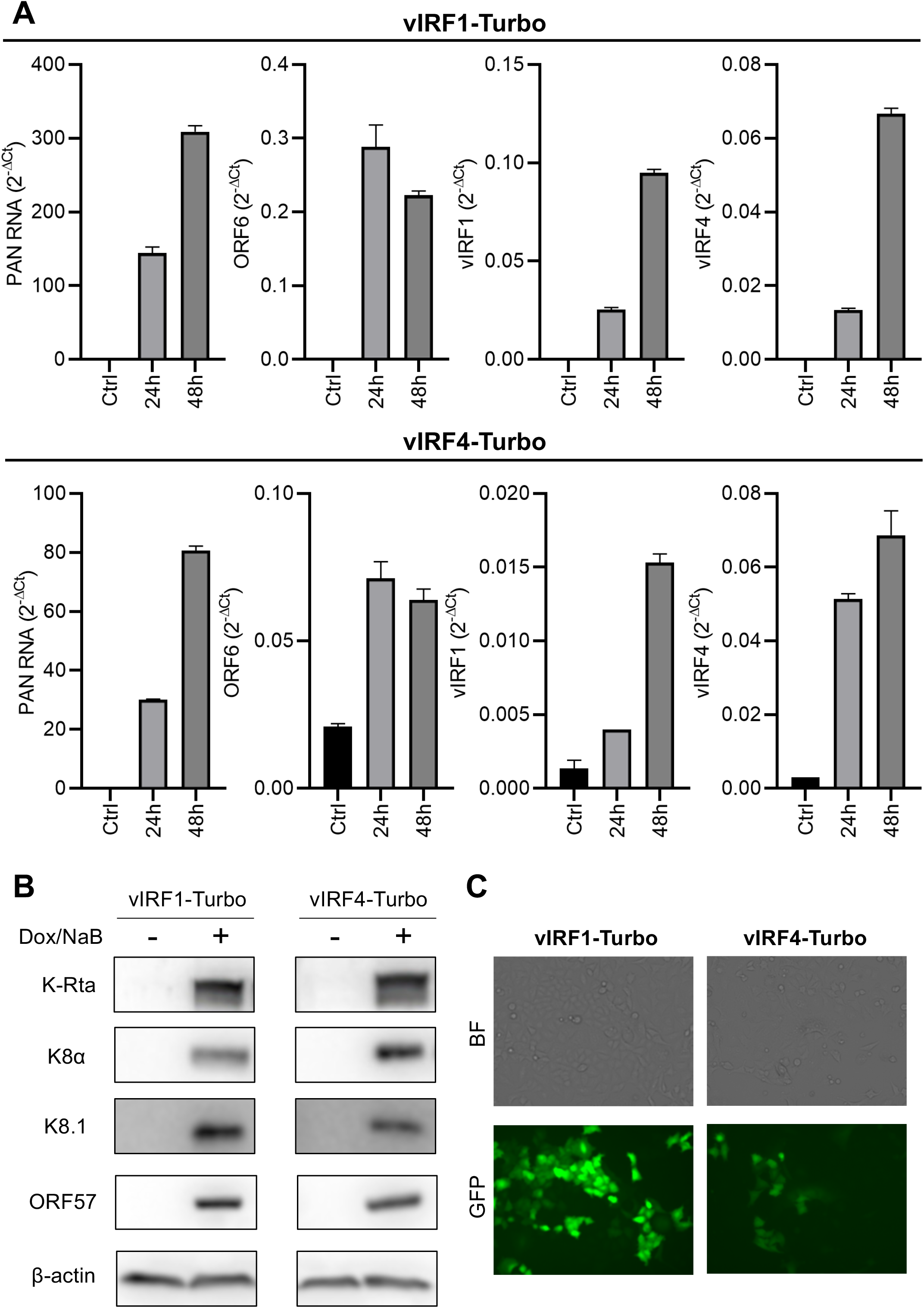
Viral gene expression and production of progeny virus. **(A) Viral gene expression for vIRF-1 and vIRF-4 mini-TurboID cells**. The vIRF-1 and vIRF-4 mini-TurboID cells were stimulated with Dox (1 μg/ml) for 24 and 48 h. Total RNA was purified at indicated time point and subjected to real-time PCR for indicated genes. Gene expression is shown as a 2^−ΔCT^. 18S ribosomal RNA was used as an internal standard for normalization. **(B) Viral protein expression in vIRF-1 and vIRF-4 mini-TurboID cells.** The vIRF-1 and vIRF-4 mini-TurboID cells were stimulated with Dox (1 μg/ml) and NaB (3 mM) for 24 h. Total cell lysates were subjected to immunoblotting using KSHV proteins and β-actin protein specific antibodies. **(C) De novo infection.** A549 cells were infected with vIRF-1 and vIRF-4 mini-TurboID virus. BF: Bright field, GFP: Green fluorescent protein.

### Proximity biotin labelling with vIRF-1 and vIRF-4

For proximity protein labeling, three replicated samples were prepared for both vIRF-1 and vIRF-4 mini-TurboID cells. Cells were reactivated with Doxycycline and NaB (sodium butyrate) for 24 h followed by addition of biotin for 1 h. Two sets of controls were also processed concurrently, in order to rule out non-specific precipitations. In the first set, the cells were left without triggering reactivation followed by incubation with biotin (+B) to rule out non-specific protein binding with biotin (Ctrl 1). For the second set, cells were reactivated with Dox/TPA for 24 h and incubated for additional 1 h in the absence of biotin (-B) to rule out non-specific interaction with streptavidin beads (Ctrl 2). Schematic workflow for the experiment is presented in **Figure 3A**. We confirmed the biotinylation signal by streptavidin blot, and vIRF-1 and vIRF-4 expression by using anti-Flag antibody (**Figure 3B**). The whole cell lysate from vIRF-1 and vIRF-4 mini-TurboID cells were further used for enrichment of biotinylated protein using magnetic beads coated with streptavidin. The enriched proteins were eluted from the streptavidin beads using trypsin on-bead digestion overnight. Ctrl1 and ctrl2 were used independently to remove background noise. We designated proteins with p-value <0.05 and fold change > 2 over both ctrl1 and ctrl2 as positive hits. Based on our setting, we identified 213 and 70 proteins from vIRF-1 and vIRF-4 mini-TurboID cells respectively (**S-Table 1, S-Table 2**).

**Figure 3.**
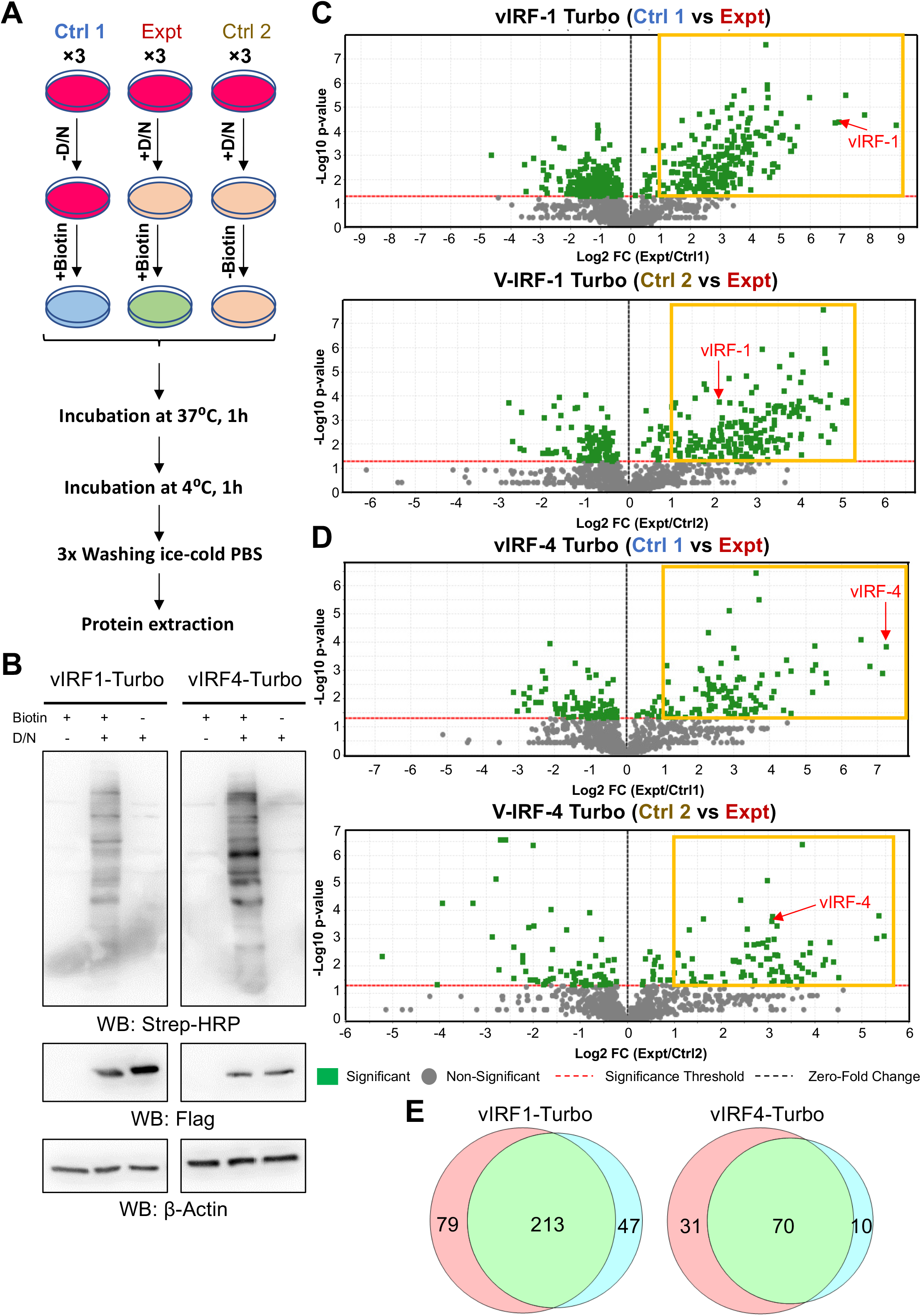
Proteins in close-proximity to vIRF-1 and vIRF-4. **(A) Schematic workflow for experimental setup.** Three biological replicates for each sample were analyzed by LC-MS/MS analysis. The plus (+) and minus (−) signs indicate presence and absence, respectively. Ctrl: Control, Expt: Experimental, D/N: Dox (1 μg/ml) and NaB (3 mM). **(B) Confirmation of biotinylation.** The cell lysate from one of the three biological replicates was subjected to immunoblotting using streptavidin HRP conjugates, Flag antibody and β-actin antibody. WB: Western Blotting. **(C-D) Identification of proteins in close proximity to vIRF-1 and vIRF-4.** Volcano plot showing differential proteins profiles in Ctrl 1 and Expt, and Ctrl 2 and Expt for vIRF-1 (**C**) and vIRF-4 mini-TurboID expressing cells (**D**). Identified and quantified biotinylated peptides are plotted as log2 fold change (Expt/Ctrl1) or (Expt/Ctrl2) versus −log10 p-value. Biotinylated peptide for vIRF-1 (**C**) and vIRF-4 (**D**) are shown with red arrow. Yellow boxes indicate selected peptides with fold change> 2 and p value< 0.05. **(E)** Venn diagram comparing proteomic lists between Ctrl1 vs Expt and Ctrl2 vs Expt (left panel for vIRF-1 and right panel for vIRF-4).

### vIRF-1 and vIRF-4 pathway analysis

Next, gene ontology (GO) analysis was performed for proteins identified in vIRF-1 and vIRF-4 mini-TurboID cells. The vIRF-1 interactome revealed significant enrichment for functions related to mRNA processing, transcription regulation by TP53, regulation of mRNA processing, and formation of RNA pol II elongation complex. Top 20 enriched GO terms are presented in **Figure 4A** (upper panel). Similarly, GO analysis for the vIRF-4 revealed again enrichment of mRNA processing, regulation of mRNA processing, mRNA polyadenylation, and mRNA splicing [**Figure 4A** (lower panel)]. Consistent with the fact that vIRF-1 and vIRF-4 have overlapping biological functions, we found overlapping possible pathway regulations in vIRF-1 and vIRF-4. Network plot by Cytoscape was generated using a subset of enriched proteins to highlight their respective protein networks (**Figure 4B**).

**Figure 4.**
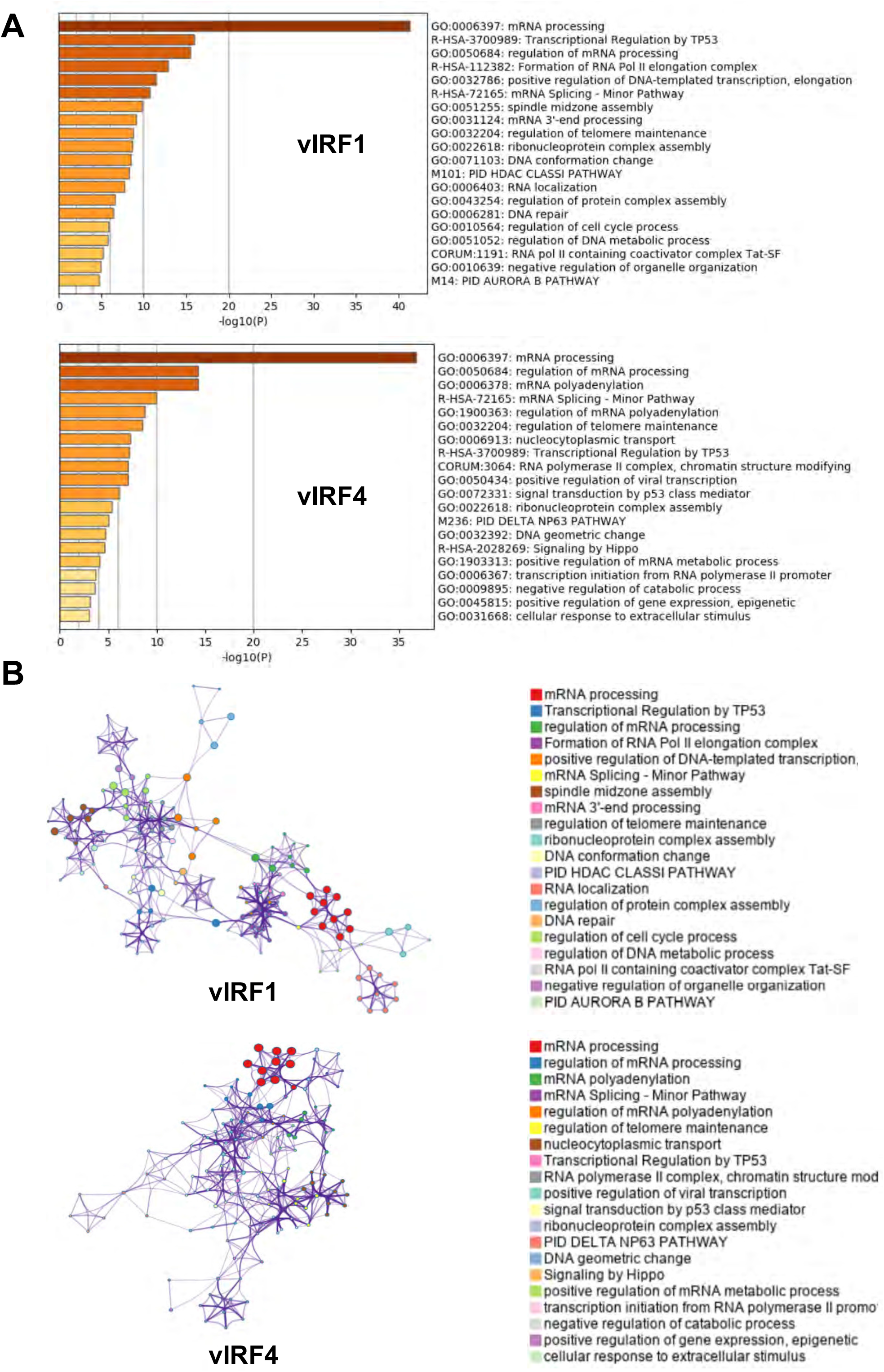
Pathway analysis for vIRF-1 and vIRF-4 interacting protein. **(A)** Top non-redundant enrichment clusters for vIRF-1 (top panel) and vIRF-4 (bottom panel) interacting proteins using Metascape bar graph (30944313). Color scales represent statistical significance. **(B)** Metascape enrichment network visualization for vIRF-1 (top panel) and vIRF-4 (bottom panel) showing the intra-cluster and inter-cluster similarities of enriched terms, up to ten terms per cluster. Cluster annotations are shown in color code.

### Effects of common hits in KSHV replication

Previous studies demonstrated that vIRF-1 and vIRF-4 possess similar biological functions to regulate interferon pathways (18, 20). We thus hypothesize that commonly targeted cellular proteins by the two viral proteins play an important role in interferon responses. Our venn diagram indicated 123 and 23 proteins were interacting exclusively with vIRF-1 and vIRF-4, respectively, and 47 proteins were found to be interacting with both vIRF-1 and vIRF-4. This suggests that the majority of vIRF-4 interacting proteins (67%) are also neighbors to vIRF-1 (**Figure 5A**). Of the 47 proteins interacting with both vIRF-1 and vIRF-4, 44 were cellular proteins whereas 3 were viral proteins (**Figure 5A**). To examine the role of those cellular proteins in KSHV replication, iSLK.219 cell line was employed. iSLK.219 carries a recombinant rKSHV.219 virus encoding a constitutively expressing GFP and an PAN RNA promoter driven RFP reporter in the viral genome, allowing us to monitor the lytic promoter activation. We used siRNA to knock-down these 44 cellular proteins followed by KSHV reactivation by treatment with Dox to induce K-Rta expression. We found that knock-down of 17 genes enhanced KSHV promoter activation, while knock-down of 6 genes lowered KSHV gene transactivation (**Figure 5B**). The corresponding images of selected knock-down experiments are shown in **Figure 5C**, and the results were further confirmed by quantifying the viral mRNAs after knock-down of selected genes, SF3B1, SF3B2 and SNW1 (**Figure 5D**). Consistent with increased viral gene expression, the viral DNA copy number in culture supernatant was increased by knocking-down of SF3B1, SF3B2 or SNW1 (**Figure 5E**). Taken together, our study suggests that some splicing factors have a role in restricting KSHV gene expression during reactivation, albeit their biological roles in general host gene transcription.

**Figure 5.**
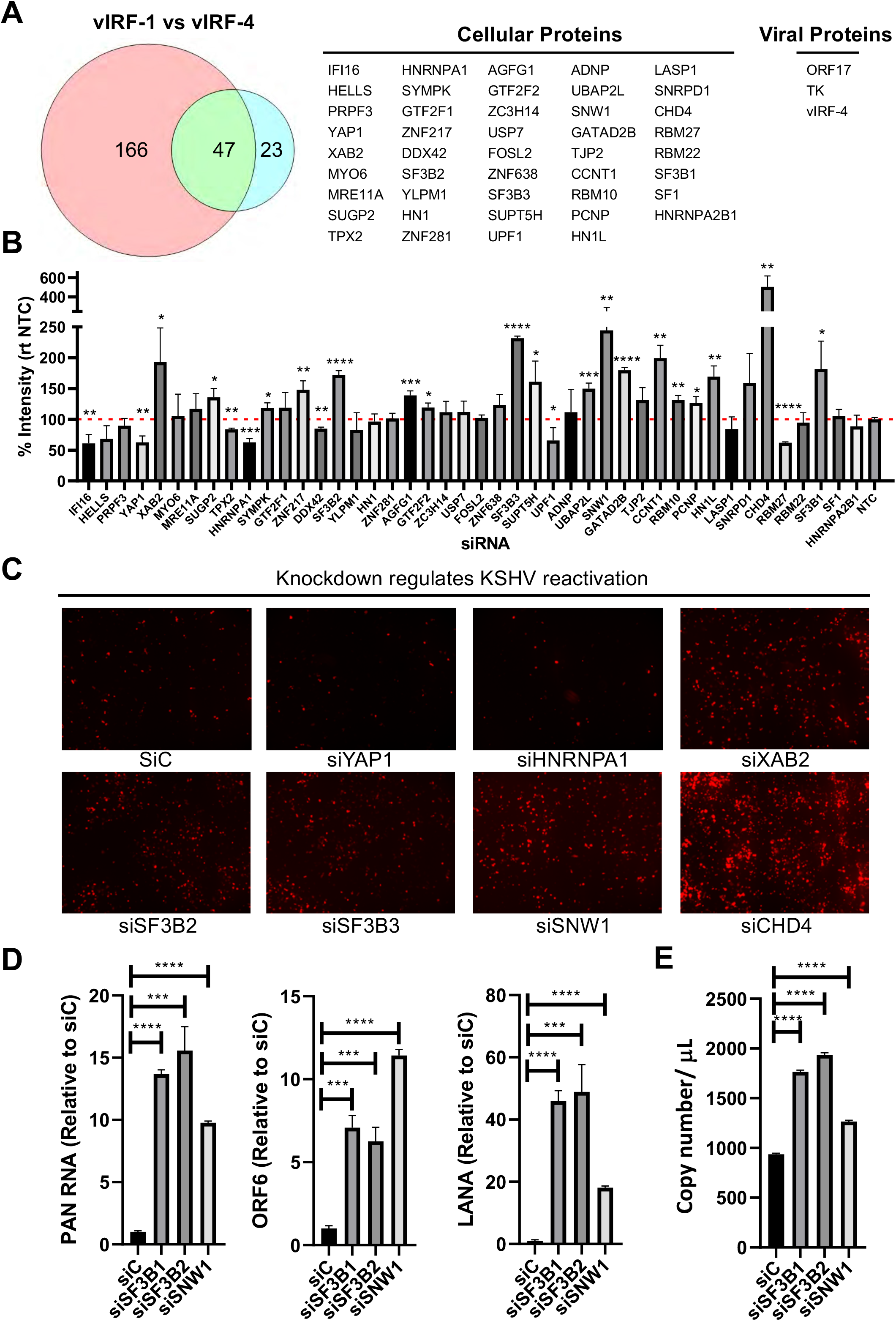
Splicing factor 3B (SF3B) subunits are suppressors for KSHV reactivation. **(A) Common protein between vIRF-1 and vIRF-4.** Venn diagram depicting proteins that interact with vIRF-1 and vIRF-4. List of cellular proteins and viral protein interacting with vIRF-1 and vIRF-4. **(B) KSHV reactivation.** Five pmol of individual siRNAs were transfected into iSLK.r219 cells for 48 h followed by reactivation with Dox (1 μg/ml) for 24 h. Percentage RFP signal was quantified relative to the non-targeting control siRNA (NTC). *p<=0.05, **p<=0.01, ***p<=0.001 and ****p<=0.0001. **(C) Microscopy imaging.** Representative RFP microscopy images of Fig 5B. **(D) Quantification of viral gene expression.** Five pmol of siC, siSF3B2 and siSNW1 were transfected in iSLK.r219 cells for 48h followed by reactivation with Dox (1 μg/ml) for 24h. PAN RNA, ORF6 and LANA gene expression was quantified using real-time PCR. ***p<=0.001 and ****p<=0.0001. **(E) Quantification of progeny virus.** Five pmol of siC, siSF3B2 and siSNW1 were transfected in iSLK.r219 cells for 48 h followed by reactivation with Dox (1 μg/ml) for 24 h. Viral copy number was quantified from tissue culture supernatant using real-time PCR. ***p<=0.001 and ****p<=0.0001.

### Splicing factor 3B (SF3B) subunits are important for IFN gene expression

Previous reports showed that the KSHV genome is sensed by RIG-I like receptors. PolyI:C is a synthetic dsRNA polymer which is recognized by RIG-I, leading to strong induction of interferons and interferon stimulatory genes (ISGs). Because KSHV vIRFs are known counteract IFN responses, we examined the relation of SF3B1 and SNW1 to interferon responses with polyI:C. The results showed that knock-down of SF3B1 or SNW1 clearly inhibited induction of type I interferon (IFNB1), type III interferon (IFNL1), and interferon downstream target gene (DDX58) [**Figure 6 (a-c)**], but not the non-IFN regulatory gene [**Figure 6(d)**].

**Figure 6.**
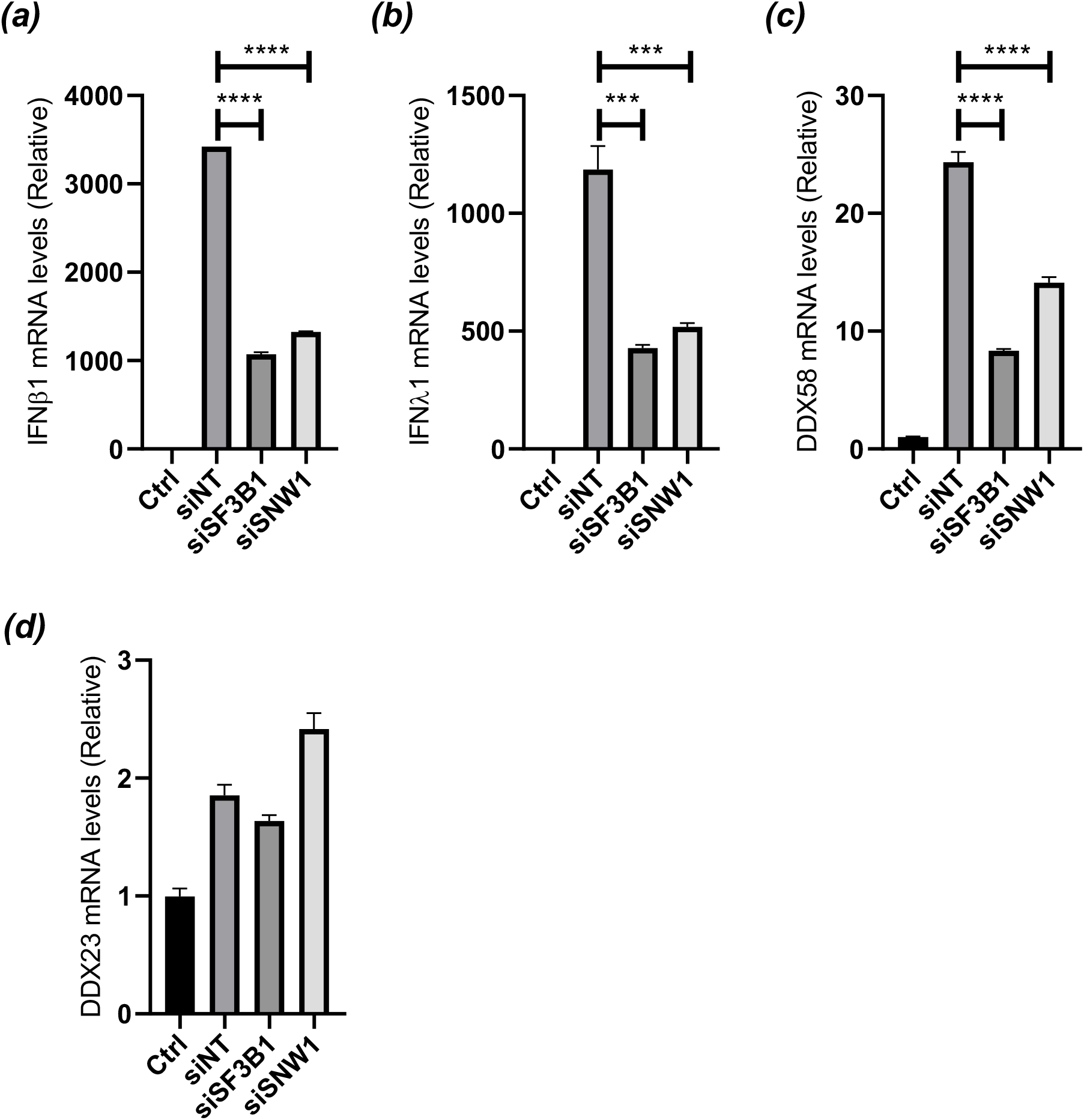
Splicing factor 3B (SF3B1) and SNW1 are suppressors for IFNB1 transcription. Five pmol of siC, siSF3B1 or siSNW1 was transfected into 293FT cells for 48 h, followed by polyI:C transfection. Twenty-four post transfection of poly:C, total RNA was harvested and measured IFN-related (a) IFNβ1, (b) IFNλ and (c) DDX58, or non-related (d) DDX23 mRNAs. ***p<=0.001 and ****p<=0.0001.

## Discussion

Since viral replication depends entirely upon host factors, understand the virus and host protein interaction network is important to find their Achilles’s heel (46). We therefore applied a mini-TurboID based system for studying the virus and host protein interaction. By constructing mini-TurboID as an integral component of KSHV BAC16 recombination system, we demonstrated a novel approach to define protein interaction networks. We propose that this approach increases the reproducibility of identifying interacting proteins, because tight interaction between biotin and streptavidin allows us to wash magnetic beads in highly stringent conditions to remove non-specific or indirect protein interactions. High reproducibility could be seen in our biological triplicated samples (S-Fig. 1).

To conveniently generate mini-TurboID tagged viruses, we first generated template plasmids similar to what we did for Rainbow-KSHV (47). With a plasmid template, homology arms can be added to primer pairs and the resultant PCR product is used for recombination (Fig. 1A). Background BAC16 can also be wild type BAC16, mutant virus, and/or Rainbow-KSHV, which allows us to examine the formation of protein complexes during viral replication and the effects of specific mutations. In this study, we used vIRF-1 and vIRF-4 as bait for validating the efficiency of PL. The vIRF-1 and vIRF-4 were selected because of their known role in regulation of innate immune response during KSHV reactivation, and multiple interacting proteins have been identified that can be used as comparisons (18, 23, 26, 48). Consistent with previous studies, vIRF-1 and vIRF-4 were found to be physically neighboring to cellular proteins that function in p53 transcriptional regulation. vIRF-1 was reported to deregulate p53 activity by interacting with ATM kinase and prevent serine 15 phosphorylation (49). In addition, vIRF-1 interacts directly with p53 to inhibit its transcriptional activation (48). Although our studies could not precipitate p53, we identified p53BP1 (p53 binding protein 1) as a possible partner of vIRF-1. We could also identify USP7 in both vIRF-1 and vIRF-4 samples, validating the PL approaches (23).

After learning that mini-Turbo worked efficiently in biotinylating cellular proteins, we tagged various other KSHV genes with mini-TurboID using the same approach. However, we learned that efficacies of biotin labeling varies significantly among different viral proteins. For example, mini-Turbo-ORF57 robustly induced biotinylated protein in total lysates with as little as 15 min of D-biotin incubation, while biotinylation by mini-Turbo-ORF50 was barely detectable in the same time frame. For this study, we also generated vIRF-2 and vIRF-3 constructs at same time; however, the level of biotinylation was lower with the same amount of D-biotin and incubation periods, leading us to drop these analyses for comparison. Differences in efficacy of biotinylation have also been seen in prior studies and abundance of viral protein expression during reactivation and subcellular nuclear localization seemed to have strong effects in the outcome of biotinylation.

Our PL studies showed a large portion of host proteins (36%) were related to mRNA processing. Within these RNA processing proteins, SF3B1, SF3B2 and SF3B3, a component of SF3b complex, were clear front runners for our further analyses. The SF3b complex is a component of the functional U2 small nuclear ribonucleoprotein (snRNP), which recognizes the exon/intron junctions and facilitates spliceosome assembly (50). Even though SF3B1 is one of many cellular genes involved in RNA splicing, *SF3B1* has been specifically identified as a commonly mutated gene in myelodysplastic syndrome (MDS) at 25–30% frequencies in MDS patients (51–53). Recent studies also showed that SF3B1 mutations increase R-loop formation and DNA damage (54). Here we found SF3B1 knock-down inhibited IFN gene expression 3 to 4-fold and also enhanced KSHV reactivation. In fact, SF3A1 and SF3B1 were reported to play a role in innate immune response to TLR ligands. The study showed that SF3A1 and SF3B1 are necessary to increase production of IL-6 and IFNβ by modulating the splicing of MyD88, an important adaptor molecule for TLR signaling pathway (55). Based on that study and ours, we propose that targeting the splicing complex might be a previously uncharacterized mechanism for KSHV to modulate host immune responses. Further studies on regulation of SF3B complex formation during KSHV reactivation and/or IFN stimulation with PL will clarify underlying mechanisms of SF3B family proteins in KSHV replication and IFN regulation.

In addition to SF3 complex, several other mRNA processing factors like XAB2, SNRPD1, SNW1, RBM10, SYMPK, and GTF2F2 were found to suppress KSHV reactivation (Fig 5). A recent study showed that SNW1 interacts with IKKγ, the regulatory subunit of IκB kinase (IKK) complex. SNW1 increases production of IL-6, IFNβ, and MX1 by enhanced activation of NF-κB and phosphorylation of TBK1 in response to influenza A virus and polyI:C (56). Influenza A virus and polyI:C are recognized by the innate immune sensor RIG-I, which plays an important role in suppressing KSHV reactivation by sensing KSHV DNA (11, 12, 57, 58). Accordingly, we explored the role of SNW1 in regulating RIG-I mediated innate immune response during KSHV reactivation. We found that knock-down SNW1 indeed enhanced KSHV replication (Fig. 5D), and this effect could be through down-regulation of IFNβ (Fig. 6).

In summary, using mini-TurboID KSHV with vIRFs as bait, we could successfully probe cellular proteins that play a role in innate immune responses. We propose mini-TurboID with recombinant KSHV BAC system as a very powerful combination to identify cellular proteins that play an important role in KSHV replication, hence a key player for respective cellular pathways.

## Acknowledgement

We would like to thank all members in Izumiya lab for valuable discussion and assistances. This research was supported by public health grants from National Cancer Institute (CA225266, CA232845), National Institute of Dental and Craniofacial (DE025985), and National Institute of Allergy and Infectious Disease (AI147207) to Y.I.

